# Integrated multi-omics analyses reveal the pro-inflammatory and pro-fibrotic pulmonary macrophage subcluster in silicosis

**DOI:** 10.1101/2024.02.16.580775

**Authors:** Hanyujie Kang, Xueqing Gu, Siyu Cao, Zhaohui Tong, Nan Song

## Abstract

**Background:** Silicosis is a lethal occupational disease caused by long-term exposure to respirable silica dust. Pulmonary macrophages play a crucial role in mediating the initiation of silicosis. However, the phenotypic and functional heterogeneities of pulmonary macrophages in silicosis have not been well-studied.

**Methods:** The silicosis mouse model was established by intratracheal administration of silica suspension. Bronchoalveolar lavage fluids (BALFs) of mice were collected for the multiplex cytokine analysis. Single-cell RNA sequencing (scRNA-seq) and spatial transcriptomics were performed to reveal the heterogeneity and spatial localization of macrophages in the lung tissues. The formation of the fibrotic nodules was characterized by histology, hydroxyproline assay, and immunohistochemical staining, respectively. The expression of the pro-inflammatory or pro-fibrotic genes was investigated by quantitative polymerase chain reaction (qPCR).

**Results:** We found that the level of pro-inflammatory cytokines and chemokines is significantly increased in the BALFs of silicosis mice. Apparent collagen deposition can also be observed in the silicotic lung tissues. By scRNA-seq, we have identified a subpopulation of Mmp12^hi^ macrophages significantly expanding in the lung tissues of mice with silicosis. Spatial transcriptomics analysis further confirmed that the Mmp12^hi^ macrophages are mainly enriched in silicosis nodules. Pseudotime trajectory showed that these Mmp12^hi^ macrophages, highly expressing both pro-inflammatory and pro-fibrotic genes are derived from Ly6c^+^ monocytes. Additionally, 4-octyl itaconate (4-OI) treatment, which can alleviate pulmonary fibrosis in silicosis mice, also reduces the enrichment of the Mmp12^hi^ macrophages. Moreover, we found a subset of macrophages in BALFs derived from patients with silicosis exhibited similar characteristics of Mmp12^hi^ macrophages in silicosis mice models.

**Conclusions:** Our study suggested that a group of Mmp12^hi^ macrophages highly express both pro-inflammatory and pro-fibrotic factors in silicosis mice, and thus may contribute to the progression of fibrosis. The findings have proposed new insights for understanding the heterogeneity of lung macrophages in silicosis, suggesting that the subset of Mmp12^hi^ macrophages may be a potential therapy target to further halt the progression of silicosis.

## 1. Introduction

Silicosis, an occupational disease with relatively high morbidity and mortality, is caused by long-term exposure to silica particles, characterized by the formation of silicotic nodules and progressive pulmonary fibrosis^1^. Recently, the incidence of silicosis gradually increases due to the development of emerging industries, such as artificial stone countertop cutting, jewelry polishing, *etc.*^2^. However, there is still no effective therapy for treating silicosis. Generally, lung transplantation is still the only treatment option for terminal silicosis. Therefore, it is urgent to explore the pathogenesis of silicosis to further find effective intervention targets.

Unlike idiopathic pulmonary fibrosis (IPF), caused by recurrent alveolar epithelial cell injury, silicosis has its unique development process^3^. Alveolar macrophages, which phagocytose silica dust, play a crucial role in the occurrence and development of silicosis^2^. Current studies indicated that an excessive inflammation triggered by silica-engulfed macrophage is the initiating factor in fibrosis^2^. Because these phagocytes cannot eliminate silica particles from the pulmonary tissue, the inflammatory damage persistently exists, followed by excessive injury repair, finally leading to the formation of silicosis nodules^3^. Thus, exploring the functions of pulmonary macrophages can help us better understand the progression of silicosis.

Conventional research on macrophage function in silicosis mainly focused on the M1/M2 polarization^4^. However, recent studies have revealed the dichotomous approach is not inadequate to present the complexity of macrophages^5^. Regarding pulmonary macrophages, these cells can be classified as alveolar macrophages (AMs) or interstitial macrophages (IMs), depending on their localization in the alveolar spaces or the lung parenchyma^6^. According to the origin, lung macrophages can be divided into resident macrophages and monocyte-derived macrophages^6^. Our group previously also explored the heterogeneity of macrophages in *Pneumocystis* -infected mice, and found a population of monocyte-derived macrophages may contribute to the clearance of infected pathogens^7^. With the development of new technologies such as single-cell-based spatial transcriptomics, the heterogeneity and spatial localization of macrophages in silicosis can be better resolved, and several subsets of macrophages have been identified as pro-fibrotic cell populations. For example, Ramachandran et al. revealed the function of TREM2^+^ macrophages in triggering fibrosis in injured liver tissue^8^. Morse et al. revealed that SPP1^hi^ macrophages contribute to lung fibrosis by activating myofibroblasts^9^. However, it is not yet known whether some specific macrophage populations are also involved in the silicosis microenvironment.

In the present study, we employed scRNA-seq, spatial transcriptomics, and multiplex cytokine analysis to discover the critical subset of pulmonary macrophages. We explored the potential contributions of disease-related macrophages in the progression of silicosis, and provided new insights into designing therapeutic therapies for silicosis by targeting specific macrophage subsets.

## 2. Methods

### 2.1. Mouse models

Eight-week-old male C57BL/6J mice were obtained from the Vital River Laboratory Animal Co., Ltd. (Beijing, China). All mice were housed in specific pathogen-free facilities. The silicosis mouse model was constructed by silica suspension administration as previously reported^10^. Briefly, each mouse was intratracheally instilled 12 mg silica (Sigma, S5631) suspended in 40 uL sterile PBS, while control mice received the same volume of sterile PBS. These mice were sacrificed in 42 days after silica exposure. The Capital Medical University Animal Care and Use Committee approved this study.

### 2.2. In vivo treatment

To establish the 4-octyl itaconate (4-OI) therapeutic model, we randomly divided mice into three groups: PBS control group, silicosis model group, and 4-OI intervention group. The method of establishing the silica-induced lung fibrosis model is the same as above. The 4-OI intervention group was administered 4-OI (50 mg/kg) (MedChemExpress, HY-112675) dissolved in sterile corn oil (MedChemExpress, HY-Y1888) by intraperitoneal injection once other days after intratracheal instillation of silica. The silicosis model group was given an equal amount of corn oil simultaneously. They were euthanized on day 42 after PBS or silica instillation.

### 2.3. Hydroxyproline assay

The contents of hydroxyproline (HYP) in the lung tissues of mice exposed to silica were measured by the Hydroxyproline Assay Kit (A030-2-1, Nanjing Jiancheng Bioengineering Institute, Nanjing, China) according to the manufacturer’s instructions. The lobe of the left lung was harvested from each mouse for detection. The absorbance of the sample was measured at 550 nm; the HYP concentration was described in the ug/left lobe.

### 2.4. Lung tissue histology

First, we fixed the mouse lung with 4% paraformaldehyde, followed by dehydrated and paraffin-embedded. Then, lung tissue sections with a thickness of 4 um were taken and incubated at 60 ℃ for 4 hours, and subsequent Hematoxylin-eosin (HE) and Masson staining were performed.

### 2.5. Immunohistochemistry

For immunohistochemistry, lung tissue sections were deparaffinized, rehydrated, and blocked endogenous peroxidase activity. The slides underwent antigen retrieval with citrate buffer pH 6.0 or tris-EDTA buffer pH 9.0. The goat serum was used in sections for blocking, followed by incubation with primary antibodies (anti-type I collagen, ABclonal; anti-Cd68, Proteintech; anti-Ly6g, ABclonal). After incubation of the secondary antibody Goat Anti-Rabbit IgG H&L (HRP) (Abcam), DAB staining and hematoxylin counterstain were done before microscopic observation.

### 2.6. Determination of cytokines levels in BALFs

Mice were sacrificed after being anesthetized with pentobarbital. Lungs underwent lavage with sterile PBS to obtain bronchoalveolar lavage fluid (BALF). After centrifugation, the supernatant was collected for the cytokines and chemokines levels measurement by ProcartaPlex Multiplex Immunoassay (ThermoFisher) based on the Luminex platform according to the manufacturer’s instructions.

### 2.7. Quantitative real-time PCR

Total RNAs were extracted from the homogenized lung tissue via TRIzol (Invitrogen Life Technologies, Carlsbad, CA, USA) and reverse-transcribed into cDNA by PrimeScript RT reagent Kit with gDNA Eraser (RR047A, Takara). The qPCR was performed with LightCycler 480 SYBR Green I Master (Roche). Then, the gene expression was calculated via the 2^−ΔΔCT^ method after normalization to β-actin. Related primers’ sequences are available on request.

### 2.8. Tissue collection for scRNA-seq and spatial sequencing

For scRNA-seq, mice were sacrificed under anesthetization and lung tissues were collected. The lung tissues were cut into fine pieces and digested at 37°C for 30 min with shaking using the Lung Dissociation Kit (Miltenyi Biotech) in combination with the gentleMACS Dissociators (Miltenyi Biotech) according to the manufacturer’s instructions. The lung homogenate was filtered with 70 µm cell strainers. Then, cell suspensions were centrifuged at 400 *g* for 6 min, and the red blood cells were removed by RBC lysing buffer. Finally, cells were centrifuged, cell counted, and viability determined.

For spatial transcriptomics, mice were euthanized and fresh lungs were harvested. The lung tissues were embedded in OCT on dry ice as quickly as possible.

### 2.9. Downloading scRNA-Seq data of human BALF

Single-cell RNA-Seq data of human BALF from two exposure miners and three patients with silicosis were downloaded from the GEO database (GSE174725).

### 2.10. ScRNA-seq and data analysis

The 10×Genomics Single Cell 3’ Library & Gel Bead Kit v3 (10×Genomics) and 10×Genomics Chromium Controller (10×Genomics) were used to generate single-cell libraries. Agilent 4200 was used to assess the quality of libraries. Then, the libraries were sequenced on the Illumina NovaSeq 6000 System (Illumina, San Diego, CA). The Cell Ranger (v.4.0.0) was applied to produce raw gene expression matrices (mouse GRCm38/mm10 using as reference genome).

Further analysis, including quality control and cell clustering, was performed in R software (v.4.2.2) using the Seurat^11^ package (v.4.1.1). For mouse scRNA-seq data if the cells were detected with less than 400 genes, more than 6,000 genes or 25000 unique molecular identifiers (UMIs), and more than 10% mitochondrial genes, then the cells were excluded from the final analysis. As for scRNA-seq data from human BALF, cells with less than 200 genes (all samples), more than 6,000 genes, or more than 10% mitochondrial genes were removed d from the final analysis. Subsequently, the NormalizeData function was used to normalize gene expression levels, and the top 2,000 variable genes were calculated by the FindVariableFeatures function. To reduce the dimensions, the scaled data was used to perform principal component analysis and t-distributed stochastic neighbor embedding (t-SNE) projections. Clusters were annotated according to the expressions of known canonical markers genes.

Differentially expressed genes and pathway enrichment

Differentially expressed genes (DEGs) were identified by the FindMarkers function. Genes expressed in > 25% of cells and with the logFC > 0.25 were selected as differentially expressed genes. The pathway enrichment analyses were implemented by the clusterProfiler R package and visualized with the ggplot2 R package. Gene Ontology gene sets were used to perform biological process (BP) analysis.

In order to compare the group 2 macrophages in human data and Mmp12^hi^ macrophages in mouse data, we produced the signatures of the two groups of macrophages based on the scRNA-seq data. The transcriptomic comparison was then performed by GSEA.

### 2.11. Scores according to inflammation and fibrosis-related gene sets

Inflammation and fibrosis-related Gene set scoring were performed using Seurat’s AddModuleScore function. The Hallmark Inflammatory Response gene set and the WP Lung Fibrosis gene set were both downloaded from the Molecular Signatures Database (http://www.gsea-msigdb.org/gsea/msigdb/index.jsp).

### 2.12. Trajectory analysis, receptor-ligand analysis, and SCENIC analysis

To infer the cell lineage developmental trajectories of macrophages and monocytes, trajectory analysis was conducted by the Monocle2^12^ R package. We used CellChat^13^ to model the receptor-ligand interaction network between cells. The pySCENIC was used for the single-cell regulatory network inference and clustering (SCENIC) analysis^14^. AUCell was used to evaluate the activity of each regulon in each cell.

### 2.13. Spatial transcriptomics data analysis

The spatial transcriptomics (ST) slide was printed with one capture area from a mouse with silicosis. Gene expression information for the ST slide was captured by the Visium Spatial platform from 10x Genomics using spatially barcoded mRNA-binding oligonucleotides in the standard protocol. Then, Space Ranger was used to quality-check and map the raw sequencing reads of spatial transcriptomics. The Seurat R package was used to analyze the gene-spot matrices produced after ST data processing from ST and the Visium sample. Spots with less than 500 genes or 1500 UMIs, and more than 25% mitochondrial genes were removed from the further analysis. The genes expressed in fewer than three spots were filtered. Normalization across spots was performed using the SCTransform function, followed by dimensionality reduction and clustering.

The multimodal intersection analysis (MIA)^15^ integrated the scRNA-seq and ST data to explore the distribution of a certain cell type in the spatial region. The MIA method is based on the hypergeometric test of the overlap between the cell type-specific genes of single-cell data and the tissue region-specific genes of ST data to assess the cell type enrichment degrees. The SpatialFeaturePlot function in Seurat generated spatial feature expression plots. DEGs of the spatial domains were identified by the FindAllMarkers function, and enrichment analysis was performed using the clusterProfiler package based on the Gene Ontology gene sets. Gene set scoring was performed using the gene set variation analysis (GSVA) package (V.1.46.0) for spatial transcriptomics data based on the gene sets in the Molecular Signatures Database.

### 2.14. Statistics

Data are presented as mean ± SD. Statistical analyses were performed by GraphPad Prism 9.0. Two groups of comparisons were evaluated by unpaired Student’s t-test. When multiple groups were compared, one-way ANOVA analysis was applied. A p-value less than 0.05 was considered statistically significant.

## 3. Results

### 3.1. Lung inflammation and fibrosis are the major hallmarks of silicosis in mouse models

To explore the complexity of cell components in the silicotic pulmonary microenvironment, we first constructed the silica-induced silicosis mouse model (Figure 1A). Hematoxylin and eosin (H&E) staining (Figure 1B) of lung tissues showed obvious silicotic nodules in the silicosis mice. Masson staining further indicated the collagen deposition in the fibrotic nodules (Figure 1B). Consistently, the measurement of hydroxyproline (HYP) contents confirmed that these silica-treated mice developed robust fibrosis 42 days post-administration (Figure 1C). The above results suggest the silica-induced lung fibrosis model exhibits the similar lung pathological features of silicotic lesions observed in patients with silicosis^1^.

**Figure 1.**
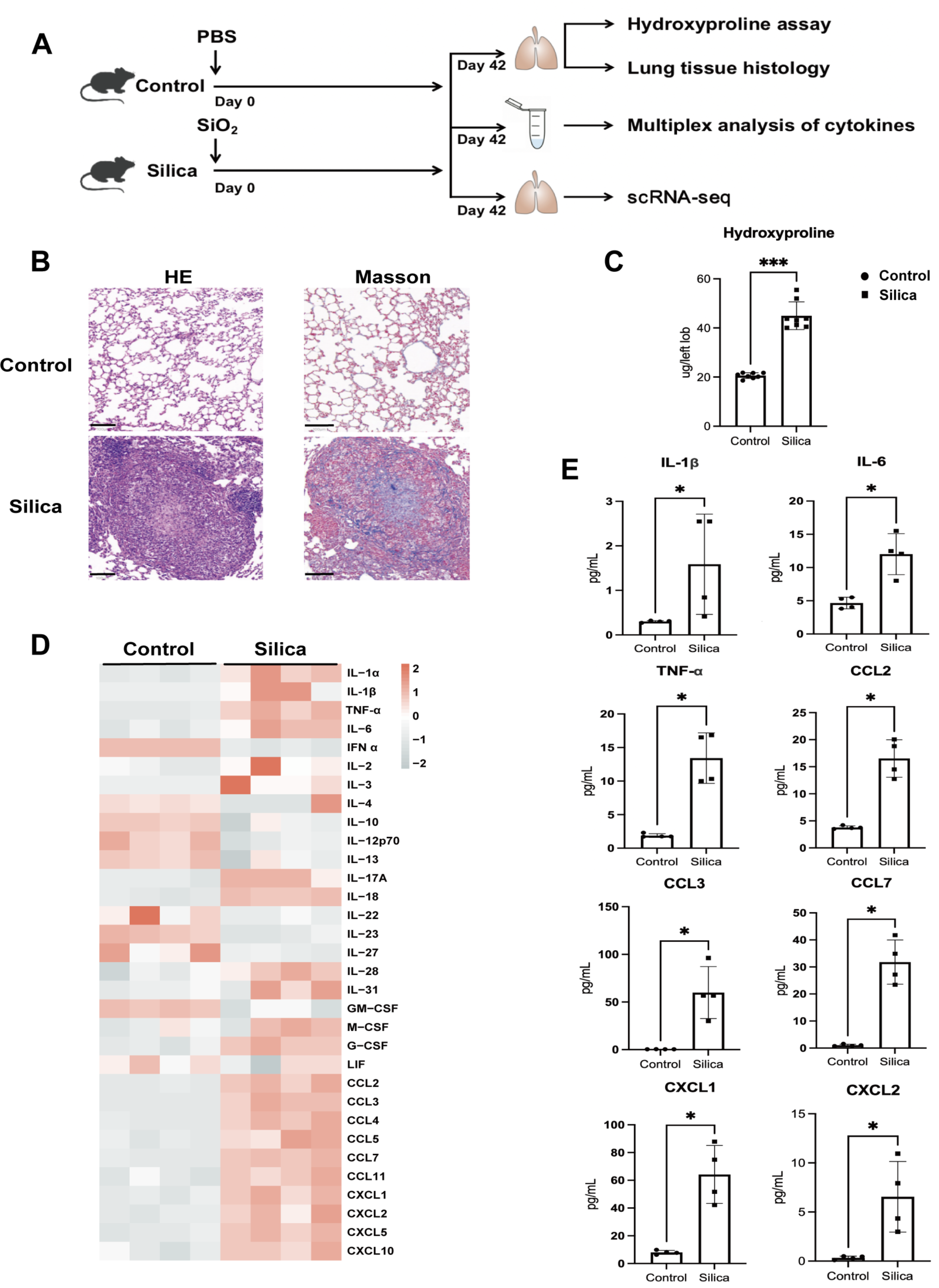
Silica-induced silicosis is characterized by significant pulmonary inflammation and fibrosis. (A) Schematic of the experimental design. (B) HE staining and Masson staining of the lung tissue of mice exposed to PBS or silica (n = 8 per group). Representative images were shown (scale bars=100 µm). (C) The hydroxyproline contents in lung tissues of mice exposed to PBS or silica (n = 8 per group). The results are shown as mean ± SD. Statistical testing was performed using an unpaired Student’s t-test. *p < 0.05, **p < 0.01, ***p < 0.001, ****p < 0.0001. (D) Heatmap for multiplex analysis of cytokines in BALFs for indicated groups of mice (n = 4 per group). (E) Bar plots showing selected differentially expressed cytokines in BALFs from (D) (n = 4 per group). The results are shown as mean ± SD. Unpaired Student’s t-test evaluated comparisons. *p < 0.05, **p < 0.01, ***p < 0.001, ****p < 0.0001.

The phagocytosis of silica particles by alveolar macrophages leads to persistent lung inflammation, which may trigger fibrosis^1^. By the quantitative polymerase chain reaction (qPCR), we found that a series of pro-inflammatory cytokines, including interleukin (IL)-1β, tumor necrosis factor-α (TNF-α), and IL-6 are markedly increased in the lung tissues of silicosis group (Supplemental Figure 1), suggesting the role of these cytokines in driving the development of lung silicosis^3^. To fully characterize the altered pro-fibrotic factors in the silicosis mice, we next determined the secretion of cytokines and chemokines in the bronchoalveolar lavage fluids (BALFs) of control and silicosis mice by cytokine multiplex analysis (Figure 1D, E). Many canonical pro-inflammatory factors, including IL-1β, IL-6, TNF-α, Ccl2, Ccl3, Ccl7, Cxcl1, Cxcl2, Cxcl10, *etc.*, were significantly elevated in mice with silicosis. Since these factors may compensate each other in triggering an excessive immune response, our results demonstrate that targeting one single factor may be insufficient for treating silicosis. We thus focus on deciphering the important cell components that contribute to the formation of the complex microenvironment.

### 3.2. Overview of Single Cells Derived from Lung Tissues of Mice with or without Silicosis

To identify the cell component(s) that may contribute to the secretion of these pro-inflammatory factors, we performed single-cell RNA sequencing (scRNA-seq) on cell suspensions of lung tissues from the silicosis mice (n=3) and normal lung samples from the control mice (n=2). After quality filtering and batch effects removal, we obtained a total of 46,006 cells for further analysis. Nine major cell clusters, including immune cells and non-immune cells, have been identified and visualized by clustering analysis (Supplemental Figure 2A, B). The distinct clusters were annotated by canonical cell type-specific genes (Supplemental Figure 2C). The atlas of immune cells demonstrated altered proportions of several cell types, including myeloid cells, neutrophils, lymphocytes, *etc.* (Supplemental Figure 2D).

We next explored the important cell components that contribute to the formation of silicosis nodules. Among the major cell clusters, the proportion of myeloid cells markedly increased in the silica exposure group, compared to those in the control group (Supplemental Figure 2D). According to the expression of marker genes (Supplemental Figure 3C), we further clustered all myeloid cells into 7 subpopulations representing macrophages, monocytes, and dendritic cells, respectively (Supplemental Figure 3A, B). Macrophages, as the most abundant population in the myeloid lineage, showed a slight increase in silicosis lung tissue (Supplemental Figure 3D). After curating the data, we found that pro-inflammatory genes (Il6, Il18, Il1b, Il1a, Tnf)^16,17^, differentially expressed chemokines (Ccl2, Ccl3, Ccl7, Cxcl1, Cxcl2, Cxcl5) detected in BALFs (Figure 1), and profibrotic genes (Il10, Mmp12, Igf1, Pdgfa, Pdgfb, Spp1, Tgfb1, Tgfbi, Lgmn, Mmp9, Mmp14, Timp1)^16,18,19^ are predominantly highly expressed in macrophages (Supplemental Figure 3E). The finding suggests that silicosis-associated pulmonary macrophages contribute to silica-induced inflammation and fibrosis.

### 3.3. Mmp12 ***^hi^*** macrophages are significantly expanded in silica-induced fibrosis lung tissue

We next explored the heterogeneity of macrophages in the silica-induced fibrotic lung tissues. The aforementioned macrophages (Supplemental Figure 3) were further divided into three clusters, including Mmp12^hi^ macrophages, Fabp4^hi^ macrophages, and proliferating macrophages, based on the differentially expressed genes (DEGs), respectively (Figure 2A-C). Among these populations, Fabp4^hi^ macrophages represent the resident alveolar macrophages (AMs), and highly express many AM-specific genes including Siglecf, Marco, Mrc1, Fabp4, Pparg, Il18, Car4, Ear2, and Ear1. Of note, the proportion of these Fabp4^hi^ macrophages within the total macrophage population in silica-exposed mice was significantly reduced, probably caused by silica particles-induced pyroptosis via activating NLRP3 inflammasome^20^ (Figure 2D). Proliferating macrophages expressed both AM-related genes (Siglecf, Marco, Mrc1, Pparg, Il18, Car4, Ear2, and Ear1) and cell proliferation signature genes (Mki67 and Top2a). The proportion of this cluster was slightly increased in silica-exposed lung tissues, indicating the recovery of the resident population following the silica-induced inflammatory response (Figure 2D). Notably, the subpopulation of Mmp12^hi^ macrophages, characterized by the high expression of the marker genes Mmp12, Fabp5, Spp1, Trem2, and Gpnmb, is predominantly enriched in silicosis lung tissues compared with normal lung tissues (Figure 2D).

**Figure 2.**
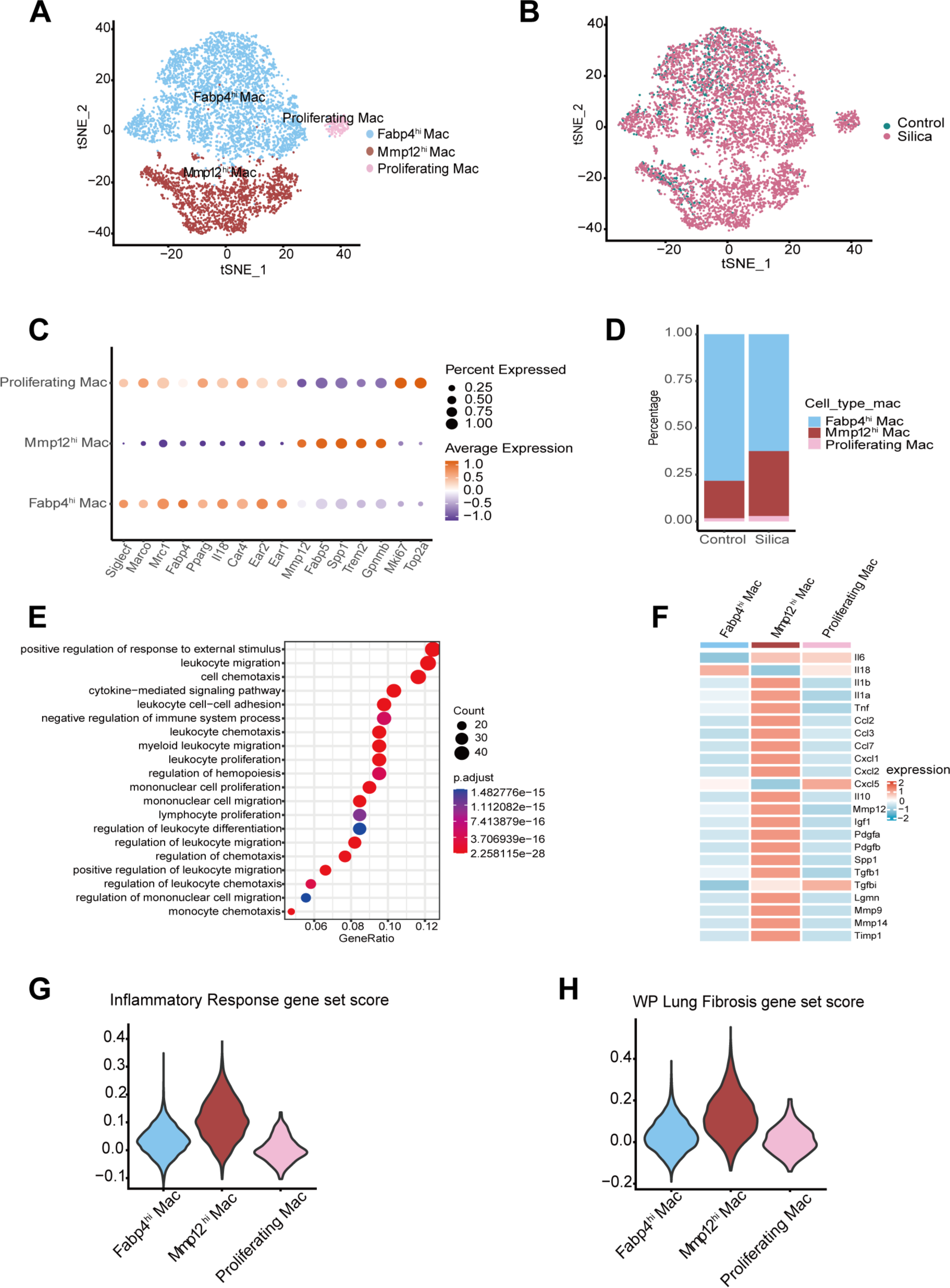
Mmp12^hi^ macrophages are enriched in silica-induced lung tissue. (A) t-SNE plot for macrophages, color-coded by cell types. (A) (B) t-SNE plot for macrophages, color-coded by groups (n = 3 in silicosis mice group and n =2 in control mice group). (B) Dot plot of the average expression of highly expressed genes for each subpopulation. The dot size shows the fraction of expressing cells, and the dots are colored based on the expression levels. (C) Bar plots showing the average proportion of each macrophage cell type from each group. (D) The GO pathway enrichment analysis using DEGs of Mmp12^hi^ macrophages versus other macrophages to explore the functions of Mmp12^hi^ macrophages. (E) Heatmap of gene expression of pro-inflammatory genes (Il6, Il18, Il1b, Il1a, Tnf), selected differentially expressed chemokines detected in BALFs in Figure 1 (Ccl2, Ccl3, Ccl7, Cxcl1, Cxcl2, Cxcl5), and profibrotic genes (Il10, Mmp12, Igf1, Pdgfa, Pdgfb, Spp1, Tgfb1, Tgfbi, Lgmn, Mmp9, Mmp14, Timp1) for each macrophage subtype from all samples. (F) Violin plot showing module scores of the Hallmark Inflammatory Response gene set in each macrophage subtype from all samples. (G) Violin plot showing module scores of the WP Lung Fibrosis gene set in each macrophage subtype from all samples.

Then, Monocle2 was utilized to infer cell trajectories for investigating the cell fate transition of these macrophage subsets (Supplemental Figure 4A). The results of pseudotime analysis showed that Mmp12^hi^ macrophages are derived from Ly6c^+^ monocytes, and then differentiated into the resident macrophages (Supplemental Figure 4A). We hypothesized that inhaled silica particles drive monocytes toward this signature of the Mmp12^hi^ macrophages. We further applied pySCENIC to assess which transcription factors underlie differences in gene expression between different macrophage subsets. We found that the transcription factor MafB is one of the top five differential transcription factors dominantly expressed in the subtype of Mmp12^hi^ macrophages (Supplemental Figure 4B). Since MafB has been known as a transcription factor that controls monocyte differentiation into monocyte-derived macrophages^21^, these analyses further supported that the Mmp12^hi^ macrophages originate from monocytes.

We then investigated the functional characteristics of Mmp12^hi^ macrophages, as these cells are significantly increased in the silicosis group. The GO pathway enrichment analysis revealed that DEGs of Mmp12^hi^ macrophages were mainly enriched in positive regulation of response to external stimulus, regulation of cell migration or chemotaxis, and cytokine-mediated signaling pathway (Figure 2E). Besides, these Mmp12^hi^ macrophages also highly expressed pro-inflammatory cytokines and chemokines, and pro-fibrotic genes (Figure 2F). Moreover, we validated these findings by using the AddmoduleScore function, and confirmed that the inflammatory response-related and lung fibrosis-related scores of Mmp12^hi^ macrophages are much higher than other subpopulations of macrophages (Figure 2G, H). These results indicated that these Mmp12^hi^ macrophages play a crucial role in the formation of silicosis nodules by triggering the pro-inflammation responses after differentiation into a pro-fibrotic phase.

### 3.4. Mmp12 ***^hi^*** macrophages are predominantly enriched in silicotic nodule lesions

Given the central role of fibroblasts in silicosis^22^, we also clustered fibroblasts into resting fibroblasts and myofibroblasts according to their characteristic genes (Supplemental Figure 5A-C). However, the proportion of myofibroblasts only showed a mild increase in the lung tissue of the silicosis group (Supplemental Figure 5D), inconsistent with previous results that the proportion of activated myofibroblasts is significantly increased in the bleomycin model of pulmonary fibrosis^23^. These data suggested that the process of digesting lung tissue for preparing single-cell suspensions may result in the loss of some specific cell populations, including myofibroblasts. To solve this problem, we further performed spatial transcriptomics to reveal the spatial localization of different types of cells in silicosis nodules.

To clarify the microenvironment of the silicotic nodules and the spatial localization of Mmp12^hi^ macrophages, we collected lung tissue from the mouse with silicosis and applied for parallel spatial transcriptomics and scRNA-seq analysis. The spatial region was first divided into two clusters by principal component analysis (PCA), consistent with the distribution of silicotic nodules and relatively normal areas in HE. Accordingly, we annotated the spatial region as “nodules of lesion” and “non-nodular area” (Figure 3A). The scRNA-seq defined similar cell populations as mentioned above (Figure 2, Supplemental Figure 2, Supplemental Figure 3).

**Figure 3.**
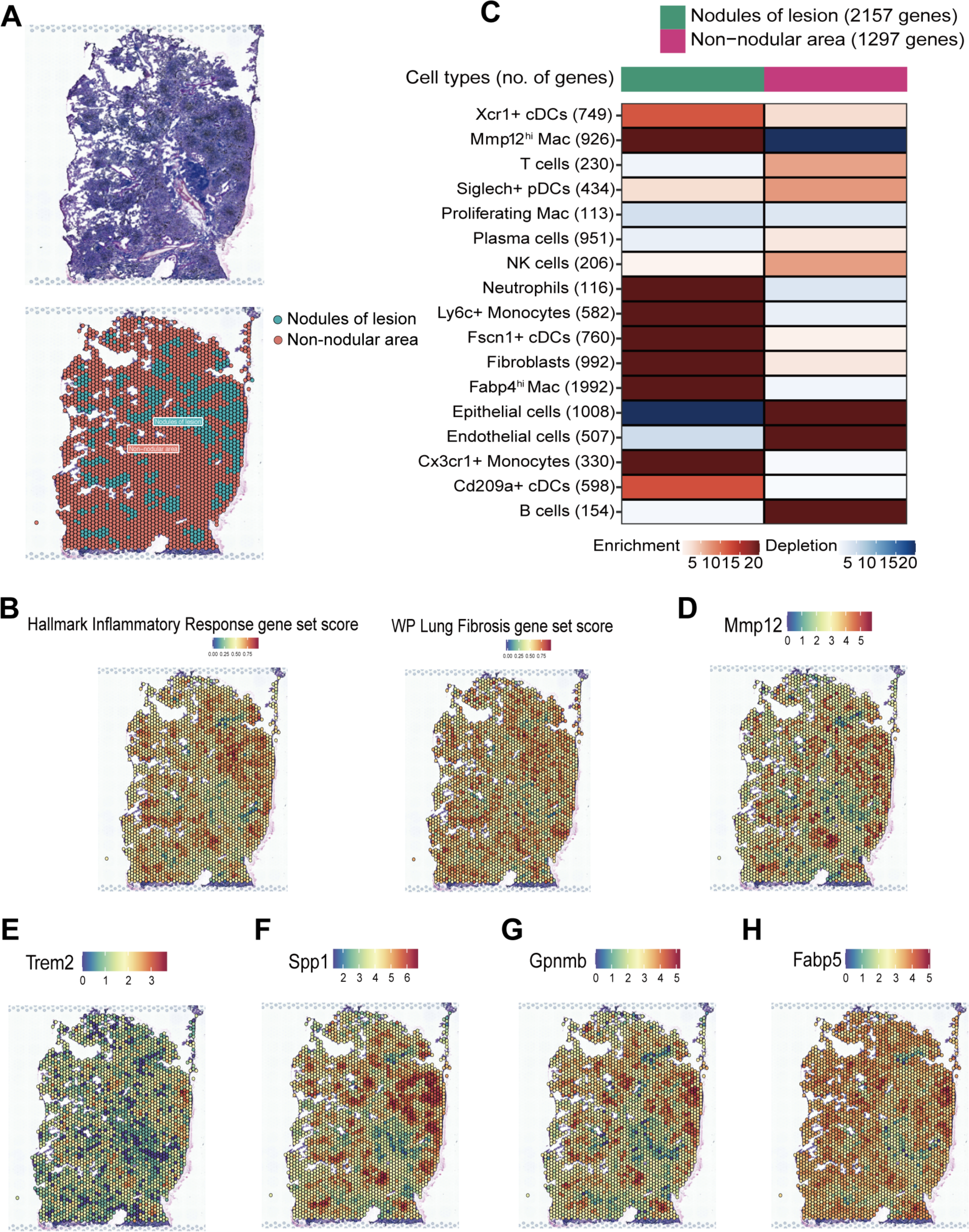
Spatial sequencing revealed the spatial localization of Mmp12^hi^ macrophages. (A) HE staining of one representative silicosis mouse lung section for spatial transcriptomics analysis (top); the spatial region was annotated as “nodules of lesion” and “non-nodular area” (bottom). (B) Spatial feature plots showing scores across lung sections of the Hallmark Inflammatory Response gene set (left) and the WP Lung Fibrosis gene set (right). (C) The silicosis MIA map of all scRNA-seq-identified cell types and ST-defined regions. The numbers of cell type- and tissue region-specific genes used in the calculation are shown. Red indicates enrichment (significantly high overlap); blue indicates depletion (significantly low overlap). (D-H) Spatially resolved gene expression of Mmp12^hi^ macrophage-associated signature genes, including Mmp12, Trem2, Spp1, Gpnmb, and Fabp5, in silicosis lung sections.

To investigate the areas of inflammation and fibrosis in the lung tissue section, we visualized module scores of the Hallmark Inflammatory Response gene set and the WP Lung Fibrosis gene set in the spatial sequencing data (Figure 3B). We found that the nodule lesion region scored highly for both inflammatory response and lung fibrosis gene sets, demonstrating higher inflammation and fibrosis activity in these regions. The results are similar to recent findings that showed a strong positive association between inflammation and fibrosis module scores in aged mice lungs post-influenza infection^24^, suggesting that the chronic inflammation is closely related with the development of fibrosis in the lung tissue with silicosis.

Previous studies have shown that inflammation influences cell-cell communications within the niche of fibrosis^25^. Thus, we performed the multimodal intersection analysis (MIA) to integrate the scRNA-seq and spatial transcriptomics data, aiming to identify the main cell population(s) enriched in the lesion niches of the lung. We found that the nodules lesion region was mainly composed of macrophages, monocytes, neutrophils, and fibroblasts (Figure 3C), confirmed by immunohistochemical staining of lung tissue from mice with silicosis for the macrophage marker Cd68, the fibroblasts marker Col1a1, and the neutrophils marker Ly6g (Supplemental Figure 5E). As expected, tissue mapping revealed that the Mmp12^hi^ macrophages localize almost exclusively to the nodule lesion area (Figure 3C). Similarly, Mmp12^hi^ macrophage-associated signature genes, such as Mmp12, Trem2, Spp1, Gpnmb, and Fabp5, are basically all relatively highly expressed in the nodule lesion region (Figure 3D-H). This result reinforced our earlier finding that these genes define a core signature for this subset of silicosis-related macrophages in the lungs. We further investigated the spatial DEGs between “nodules of lesion” and “non-nodular area”. The GO pathway enrichment analysis suggested that many enriched pathways, such as positive regulation of response to external stimulus, cell chemotaxis, regulation of leukocyte migration, and cytokine-mediated signaling pathway, in the nodules lesion region are similar to those found previously in Mmp12^hi^ macrophages (Supplemental Figure 5F). In conclusion, these results suggested that Mmp12^hi^ macrophages are mainly enriched in the silicotic nodules region, and may contribute to promoting the formation of pro-fibrotic microenvironment.

### 3.5. Cell-cell interactions between Mmp12 *^hi^* macrophages with neutrophils and fibroblasts may play a critical role in silicosis

To further infer the role of Mmp12^hi^ macrophages in fibrotic nodules, we used the ligand-receptor interaction tool CellChat to identify potential cell-cell communication as described^13^. Circle plots showed the number and strength of outgoing signals from Mmp12^hi^ macrophages to neutrophils or fibroblasts, both increased in the silicosis group compared with the control group (Figure 4A, B). Information flow analysis showed that several signaling pathways associated with pro-fibrosis activity, such as FN1 and SPP1, are upregulated under silicosis conditions (Figure 4C). Signaling pathways involving cytokines and chemokines, including CCLs and CXCLs, were also markedly upregulated in silica-induced lung tissue. We further inferred the ligand-receptor interactions between macrophages and other cells (monocytes, neutrophils, and fibroblasts) in silicosis and normal lung tissues. In particular, the ligand-receptor interactions related to cytokine and chemokine pathways between Mmp12^hi^ macrophages and neutrophils are significantly enhanced in the silicosis model group (Figure 4D). Of note, the ligand-receptor interactions associated with pro-fibrosis, such as SPP1-CD44 and FN1-ITGAV/ITGB1, which are absent between Mmp12^hi^ macrophages and fibroblasts in the control group, are highly activated in silicosis group (Figure 4E). Taken together, these results indicate the interactions of Mmp12^hi^ macrophages with neutrophils and fibroblasts, are critical event that modulates the inflammatory damage and shapes a pro-fibrotic environment.

### 3.6. Group 2 macrophages in silicosis patients have similar gene expression features to Mmp12 ***^hi^*** macrophages in mice

To further validate the above findings in human samples, we reanalyzed published scRNA-seq data for the BALFs derived from three patients with silicosis and two co-workers without silicosis^26^. A total of 22601 cells from five samples were obtained for subsequent analysis. Based on the canonical signature genes, we identified several major clusters including myeloid cells, neutrophils, T cells, B cells, megakaryocytes, and epithelial cells (Supplemental Figure 6A, B).

Macrophages were further extracted from the myeloid lineage (Supplemental Figure 6C, D) and re-clustered to evaluate the heterogeneity. According to the characteristic genes, we divided these macrophages into three groups, including group 1 macrophages (Mac.Group1), group 2 macrophages (Mac.Group2), and proliferative macrophages, respectively (Figure 5A-C). We found that similar to Mmp12^hi^ macrophages in mice, the group 2 macrophage in the BALFs of human subjects exhibited characteristic signatures including MMP12, SPP1, FABP5, GPNMB, and TREM2. Moreover, the group 2 macrophages are also significantly enriched in the BALFs of silicosis patients (Figure 5D). The GO pathway enrichment analysis of DEGs of group 2 macrophages revealed that pathways enriched in these cells highly resemble those of Mmp12^hi^ macrophages in mice (Supplemental Figure 6E). By comparing the transcriptomic features, we found that human group 2 macrophages show similar signatures with Mmp12^hi^ macrophages in mice, and express high levels of pro-inflammatory and pro-fibrotic related genes (Figure 5E-G), suggesting the functional similarity of the subset of macrophages in human and mice. These results suggested the potential pro-inflammatory and pro-fibrotic role of the group 2 macrophages in the pathogenesis of silicosis patients, which is consistent with the role of Mmp12^hi^ macrophages in mice.

**Figure 4.**
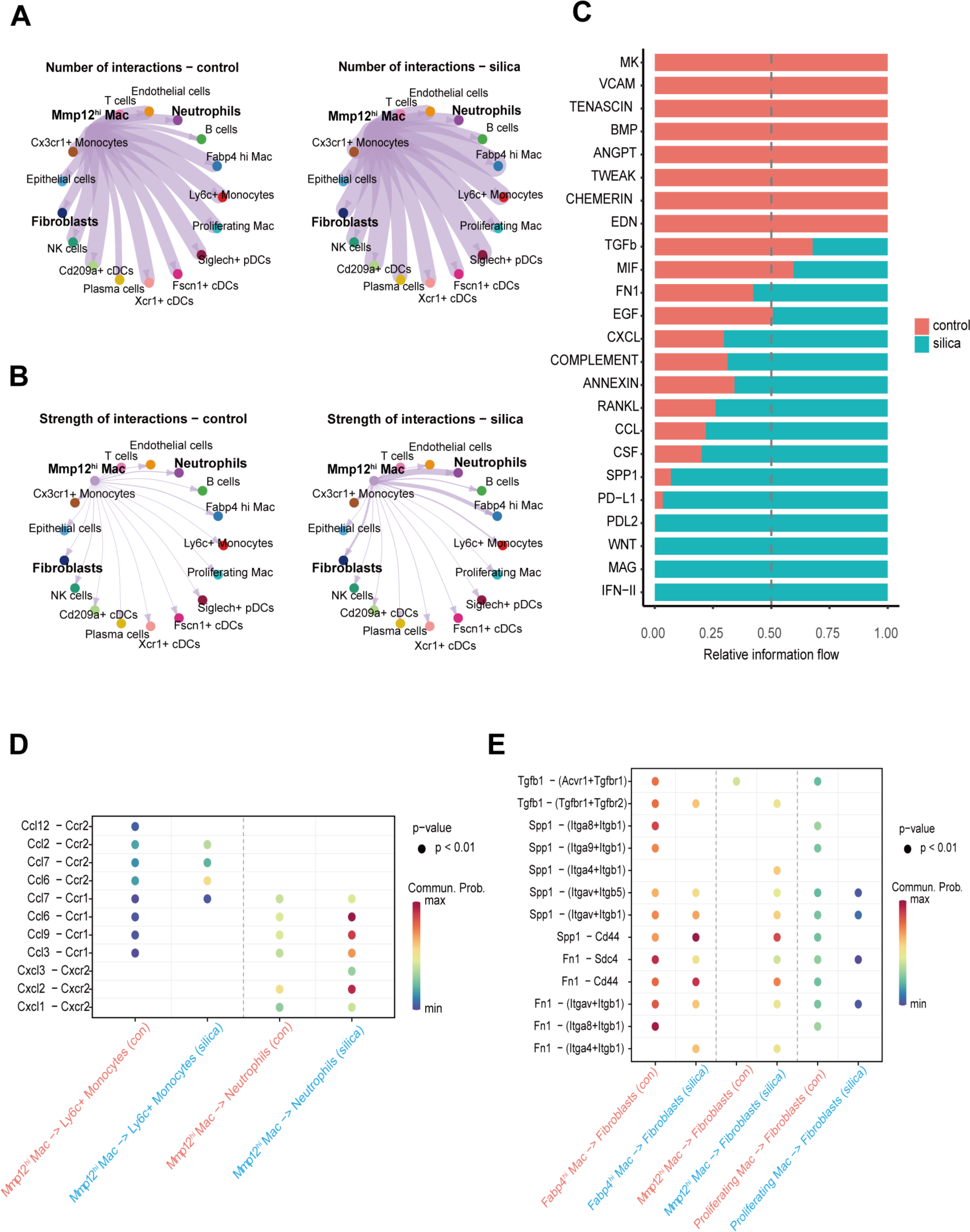
Cellchat analysis of the communication between cells in the control and silicosis group. (A) Visualization and analysis of cell-cell communication in control and silicosis group using CellChat. Circle plots placing Mmp12^hi^ macrophage subsets as the central nodes of analysis. An interaction between a pair of cell types is depicted by a line connecting two cell types. The thickness of the line represents the number of that interaction. (B) Visualization and analysis of cell-cell communication in control and silicosis group using CellChat. Circle plots placing Mmp12^hi^ macrophage subsets as the central nodes of analysis. An interaction between a pair of cell types is depicted by a line connecting two cell types. The thickness of the line represents the strength of that interaction. (C) Signaling pathways ranked by differential overall information flow of inferred interactions in control (red) and silicosis (blue) samples. (D) Dot plot showing predicted interactions between CCL and CXCL chemokine ligands produced by Mmp12^hi^ macrophages and receptors on neutrophils or monocytes at the control or silicosis mice group. Color denotes communication probability; size denotes p-value. (E) Dot plot showing predicted interactions between fibrosis-related FN1, SPP1, and TGFb ligands produced by Mmp12^hi^ macrophages and receptors on fibroblasts at the control or silicosis mice group. Color denotes communication probability; size denotes p-value.

**Figure 5.**
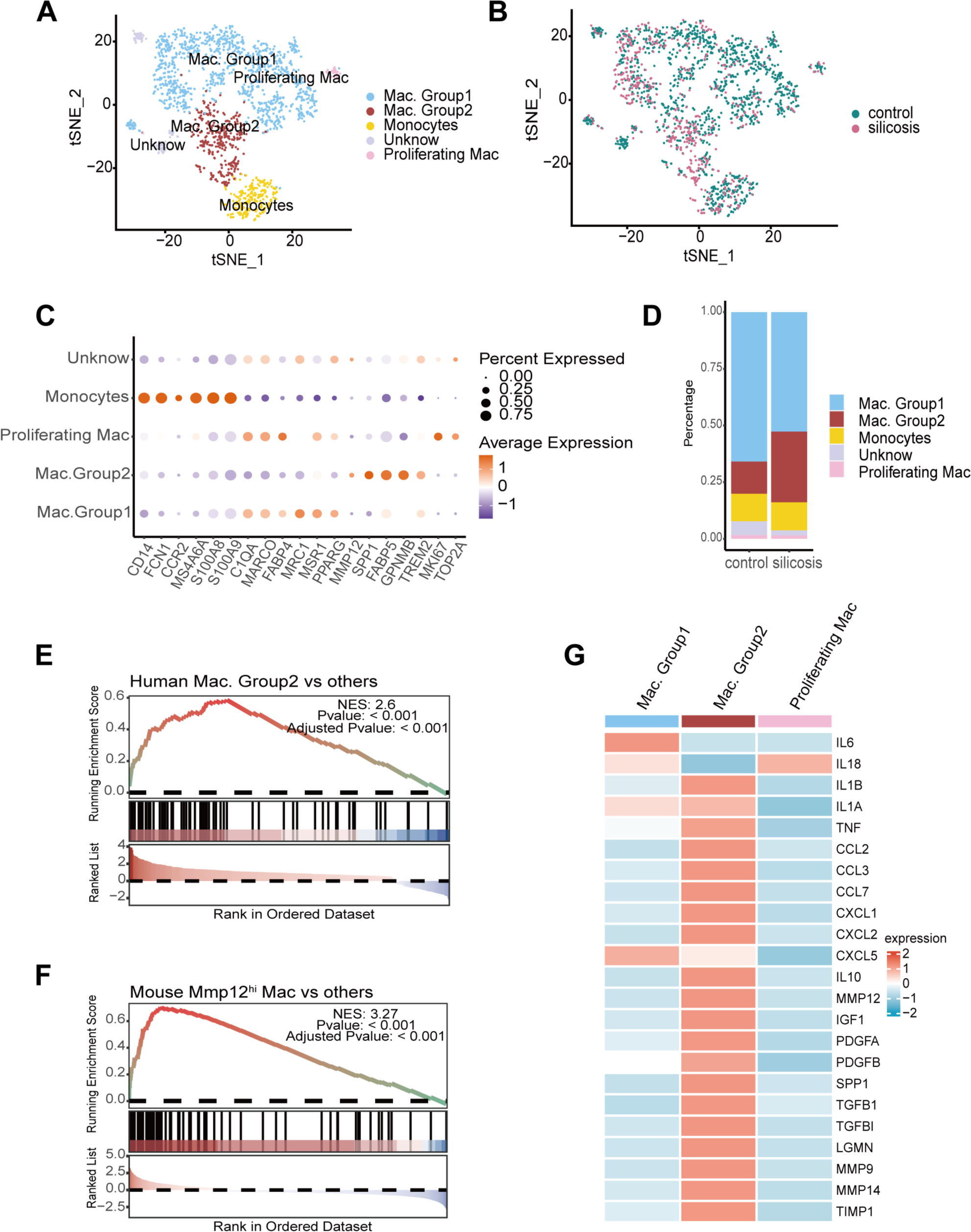
Group 2 macrophages in patients with silicosis resemble Mmp12^hi^ macrophages in silica-induced mice models. (A) t-SNE plot for macrophages and monocytes, color-coded by cell types. (B) t-SNE plot for macrophages and monocytes, color-coded by groups (n = 3 in silicosis patients group and n =2 in control group of exposure miners). (C) Dot plot of the average expression of highly expressed genes for each subpopulation. The dot size shows the fraction of expressing cells, and the dots are colored based on the expression levels. (D) Bar plots showing the average proportion of each cell type from each group. (E) GSEA analysis of DEGs of human Group 2 macrophages versus other macrophages using signatures of Mmp12^hi^ macrophages in mouse dataset. (F) GSEA analysis of DEGs of mouse Mmp12^hi^ macrophages versus other macrophages using signatures of Group 2 macrophages in human dataset. (G) Heatmap of gene expression of pro-inflammatory genes (IL6, IL18, IL1B, IL1A, TNF), selected differentially expressed chemokines detected in BALFs in Figure 1 (CCL2, CCL3, CCL7, CXCL1, CXCL2, CXCL5), and profibrotic genes (IL10, MMP12, IGF1, PDGFA, PDGFB, SPP1, TGFB1, TGFBI, LGMN, MMP9, MMP14, TIMP1) for each macrophage subtype from all samples.

### 3.7. 4–OI may ameliorate silicosis-related inflammation and lung fibrosis by affecting Mmp12^hi^ macrophages

In addition to the expression of Mmp12, Spp1, Gpnmb, Fabp5, and Trem2, immune responsive gene 1 (Irg1) is also markedly increased in the Mmp12^hi^ macrophages of silicosis mice (Figure 6A). Irg1, highly expressed in activated macrophages, can mediate itaconate production in response to inflammation^27^. Ogger et al. have reported the role of itaconate in controlling the severity of pulmonary fibrosis^28^. As the cell-permeable derivative of itaconate, 4-OI has been widely investigated due to the anti-inflammatory activity^29,30^. The anti-fibrotic effect of 4-OI has also been documented in the bleomycin-induced pulmonary fibrosis model^31^. Some studies have revealed that 4-OI can significantly reduce macrophage infiltration in lung tissue during inflammatory diseases^32,33^. We thus designed *in vivo* studies to determine the therapeutic effect of 4-OI in silicosis.

**Figure 6.**
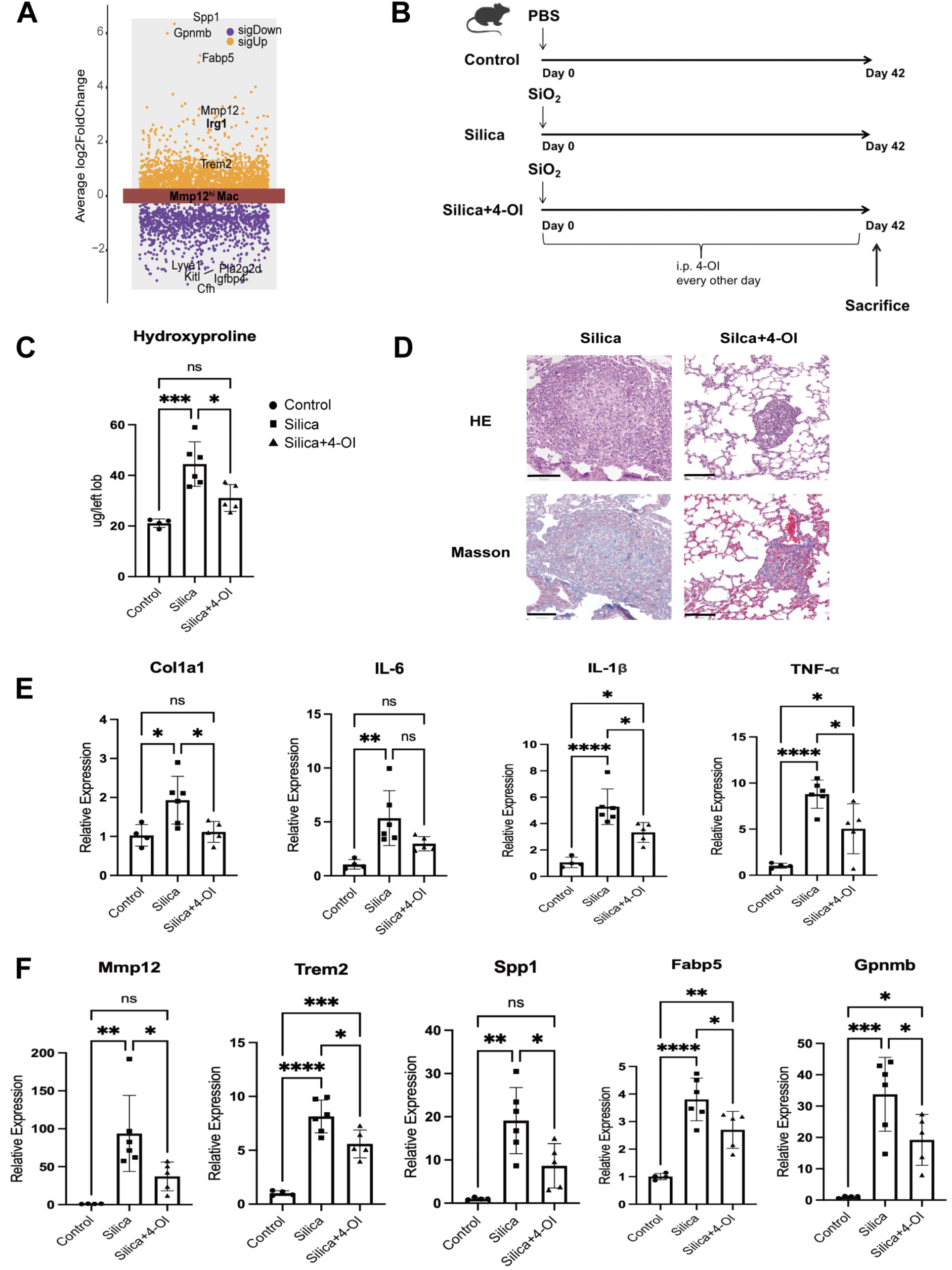
4-OI treatment may partially rescue the pulmonary inflammation and fibrosis in mice with silicosis by reducing the subset of Mmp12^hi^ macrophages. (A) Volcano plot showing the DEGs for Mmp12^hi^ macrophages in silicosis mice versus control mice. (B) Schematic plot for the 4-OI treatment schedule and study design. (C) The hydroxyproline contents in mice lung tissues of different groups (n = 4-6 per group). The results are shown as mean ± SD. One-way ANOVA evaluated comparisons for multiple comparisons. *p < 0.05, **p < 0.01, ***p < 0.001, ****p < 0.0001. (D) HE staining and Masson staining of the mice lung tissues of different groups (n = 4-6 per group). Representative images were shown (scale bars=100 µm). (E) mRNA expression of Col1a1, Il-1β, Tnf-α, and Il6 in mice lung tissues from the indicated group of mice in (B) was detected by qPCR (n = 4-6 per group). The results are shown as mean ± SD. One-way ANOVA evaluated comparisons for multiple comparisons. *p < 0.05, **p < 0.01, ***p < 0.001, ****p < 0.0001. (F) mRNA expression of Mmp12^hi^ macrophage-associated signature genes in mice lung tissues from the indicated group of mice in (B) was detected by qPCR (n = 4-6 per group). The results are shown as mean ± SD. One-way ANOVA evaluated comparisons for multiple comparisons. *p < 0.05, **p < 0.01, ***p < 0.001, ****p < 0.0001.

Mice were intratracheally administered sterile PBS or silica suspension at day 0. Solvent or 4-OI was intraperitoneally injected every other day for treatments (Figure 6B). Silicosis mice were sacrificed at 42 days after PBS or silica instillation, and analyzed with histology and hydroxyproline detection kit (Figure 6B-D). We found silicosis-related pathological changes in H&E staining are obviously alleviated after 4-OI treatment (Figure 6D). Moreover, 4-OI therapeutically inhibited the mRNA expression levels of pro-inflammatory factors, including IL-1β, IL-6, and TNF-α, in lung tissues (Figure 6E). The severity of pulmonary fibrosis, as evaluated by the Masson staining, the hydroxyproline test, and Col1a1 mRNA levels, was also attenuated by the 4-OI treatment (Figure 6C-E). More importantly, the mRNA expression levels of characteristic genes associated with Mmp12^hi^ macrophages in lung tissues were significantly reduced in the 4-OI treatment group (Figure 6F). Taken together, our results indicated that 4-OI may attenuate the lung inflammatory response and fibrosis in the silicosis mouse model by reducing the infiltration of Mmp12^hi^ macrophages. Targeting these Mmp12^hi^ macrophages is expected to become a potential way to treat silicosis.

## 4. Discussion

Silicosis is a potentially fatal occupational disease characterized by lung inflammation, pulmonary fibrosis, and impaired lung function, but current therapeutic strategies are limited, probably due to unclear molecular mechanism^1^. Though activated fibroblasts are the major component in fibrotic lesions, the role of macrophages has been highlighted in inflammation, tissue remodeling, and fibrosis^6^. Therefore, it is essential to identify disease-associated macrophage subpopulations in silicosis to guide precision therapy. In this study, we used multiplex cytokine analysis, scRNA-seq, and spatial transcriptomics to reveal the silicosis microenvironment. We found the levels of many canonical pro-inflammatory cytokines and chemokines were significantly increased in the BALFs derived from mice with silicosis. We further identified a population of monocytes-derived Mmp12^hi^ macrophages that expand markedly in silica-induced fibrosis lung tissue. These Mmp12^hi^ macrophages are mainly localized to silicotic nodule lesions and highly express inflammation-related cytokines, chemokines, and fibrosis-related factors, suggesting a potentially crucial role in silicosis. Mmp12^hi^ macrophages in the mouse model shared similar gene expression features and functions with the subsets in BALFs of patients with silicosis. These findings provide possible targets for clinical intervention of silicosis.

Pulmonary macrophages are heterogeneous populations that can be divided into subsets based on their localization, origin, function, *etc.*^6^. Lv et al. have found that in bleomycin-induced pulmonary fibrosis, macrophage subpopulations change dynamically with disease progression^34^. The steady-state AMs quickly decreased during inflammatory responses and replenished with monocyte-derived macrophages^34^. Misharin et al. have revealed the contribution of monocyte-derived macrophages on the pathogenesis of lung fibrosis^35^. Our study found that a population of monocyte-derived macrophages characterized by relatively high expression of Mmp12 is significantly enriched in silica-induced lung fibrosis, suggesting these cells may promote the pathogenesis of silicosis. Yi et al. have found that MMP12 may contribute to the development of subretinal fibrosis by promoting macrophage-to-myofibroblast transition^36^. In the present study, we focused on the characteristics of this subpopulation of Mmp12^hi^ macrophages, other than Mmp12 itself. Our scRNA-seq data revealed that this subset, besides highly expressing Mmp12, also expresses a series of characteristic genes, such as Spp1, Gpnmb, Fabp5, and Trem2. These silicosis-related genes observed in Mmp12^hi^ macrophages were also found highly expressed in some subsets of macrophages found in other diseases, such as scar-associated pro-fibrotic TREM2^+^ CD9^+^ macrophages in liver fibrosis^8^ and lipid-associated macrophages in obese adipose tissue^37^.

Our study indicated that silica-induced pulmonary fibrosis was characterized by the depletion of Fabp4^hi^ AMs and the enrichment of monocyte-derived Mmp12^hi^ macrophages in the lung. Wang et al. also found that proinflammatory monocyte-derived interstitial macrophage induced by silica contributed to the pathogenesis of silicosis^38^. Our results showed that the DEGs of Mmp12^hi^ macrophages were enriched in positive regulation of response to external stimulus, regulation of cell migration or chemotaxis, and cytokine-mediated signaling pathway. To some extent, these were consistent with the results of multiplex cytokine analysis of BALFs from mice, which suggested an enrichment of inflammation-related cytokines and chemokines in the silicosis microenvironment. These findings indicated the critical role of Mmp12^hi^ macrophages in guiding the migration of other inflammatory cells by producing various cytokines or chemokines when exposed to external stimuli, which may contribute to silica-induced lung inflammation injury. Our previous study revealed that Mmp12^+^ macrophages were the primary source of inflammatory factors during *Pneumocystis* infection^7^. Conway et al. identified a novel cluster of Mmp12^+^ macrophages that play the role of repair after the kidney injury, and they resemble the transcriptome of monocytes that infiltrate the kidney during recovery from acute ischemia-reperfusion injury^39^. Consistent with these studies, we found that silicosis-related Mmp12^hi^ macrophages highly expressed pro-inflammatory and pro-fibrotic genes. The inflammatory response-related and lung fibrosis-related scores were also highest in Mmp12^hi^ macrophages among all subpopulations of macrophages, demonstrating that Mmp12^hi^ macrophages, exhibited a mixed M1/M2 phenotype, contributing to both inflammation and fibrosis in silicosis.

Silicotic nodules are composed of two distinct zones, termed as central and peripheral zone, respectively. The central zone contains collagen fibers, while the peripheral zone mainly consists of fibroblasts and dust-laden macrophages^1^. We used spatial transcriptomics combined with scRNA-seq to characterize the distribution of cell subtypes in silicosis tissues and the microenvironment of silicotic nodules. Our results similarly found that the predominant components of silicotic nodules lesions were macrophages, monocytes, neutrophils, and fibroblasts. Macrophages are the most common type of immune cell in silicotic nodules. Importantly, we found that this subpopulation of Mmp12^hi^ macrophages, which expand markedly in silica-induced fibrosis lung tissue, are distributed almost exclusively in the region of silicotic nodule lesions. Besides, the cell-cell interaction circle plots suggest the increased number and intensity of interactions between this subset of macrophages with other cells distributed in silicotic nodules, such as neutrophils and fibroblasts. On the one hand, monocyte-derived Mmp12^hi^ macrophages may secrete several pro-inflammatory and pro-fibrotic factors that contribute to a microenvironment of persistent inflammation and fibrosis in silicotic nodules. On the other hand, the cluster of Mmp12^hi^ macrophages activates neutrophils or fibroblasts directly. Crosstalk between macrophages and neutrophils or fibroblasts has been widely reported in the tumor microenvironment^40,41^, and disrupting these interactions has been considered a potential treatment target to improve immunotherapy. In summary, our results suggested a central role for Mmp12^hi^ macrophages in the cell interaction network in the silicosis microenvironment. The interactions of Mmp12^hi^ macrophages with neutrophils and fibroblasts in silicotic nodules still need to be deeply studied in the future, and exploring the method to block the interaction between Mmp12^hi^ macrophages and other cells may be a novel strategy for silicosis treatment.

4-OI is a cell-permeable derivative of itaconate, which has received much attention for its ability to exert antioxidant and anti-inflammatory effects through activating nuclear factor-related factor 2 (Nrf2)^29^. Prior studies have demonstrated the ability of 4-OI to inhibit lung macrophage infiltration in inflammatory diseases^32,33^. However, the effectiveness of 4-OI against silicosis is unknown. Therefore, we sought to explore the therapeutic roles of 4-OI in silicosis mice models. Our results showed that 4-OI could ameliorate silica-induced pulmonary inflammation and fibrosis. We further found that 4-OI significantly decreased the mRNA expression levels of Mmp12^hi^ macrophages-related genes in lung tissues, implying the reduction in the infiltration of Mmp12^hi^ macrophages. These data provided a potential explanation for the protective role of 4-OI in silicosis and suggested that 4-OI may ameliorate silicosis by blocking the recruitment of disease-related monocytes-derived Mmp12^hi^ macrophages to lung tissues.

Similar to Mmp12^hi^ macrophages, a subset of pro-fibrotic macrophages is also enriched in BALF of silicosis patients (Figure 5), highly resembling the pro-fibrotic SPP1^hi^ macrophages found in the lungs of IPF patients^9^. Park et al. reported that disease-associated macrophages are potential targets for intervention^42^. Nowadays, some studies about therapeutic strategies targeting macrophages in pulmonary fibrosis focus more on modulating macrophage polarization^43,44^. It has been explored that the effect of specifically depleting a specific subpopulation of macrophages in BLM-induced pulmonary fibrosis^45^, with the finding that more collagen deposition and inflammatory cell infiltration were observed in the lungs of mice lacking the specific macrophage subset. In silica-induced pulmonary fibrosis models, depletion of pro-fibrotic C1q^+^ interstitial macrophages has been found to inhibit fibroblast activation but promote inflammation^19^. Therefore, targeted macrophage therapy still has a long way to go, and our findings may provide some reference value for targeted macrophage therapy in silicosis.

In conclusion, we first proposed a subset of pro-fibrotic Mmp12^hi^ macrophages featured by conferring considerable pro-inflammatory and pro-fibrotic effects in the lungs of mice with silicosis by scRNA-seq analysis. We also clarified the localization of Mmp12^hi^ macrophages and their crosstalk with other neighboring cells by spatial transcriptomics and cell-cell interaction analysis, revealing their potential roles in silicotic nodule lesions formation. Treatment of 4-OI exerted anti-inflammatory and anti-fibrotic effects in silicosis, probably by reducing the recruitment of Mmp12^hi^ macrophages. These findings illustrated that Mmp12^hi^ macrophages may represent potential targets for treating silicosis individuals.

## Data availability statements

The data generated during the current study are available from the corresponding author on reasonable request.

## Ethics Statement

The animal study was approved by the Animal Experiments and Experimental Animal Welfare Committee of Capital Medical University.

## Disclosure

These authors report no conflicts of interest in this work.

## Author contributions

ZT and NS conceived and designed the study. HK, XG, and SC performed the experiments. HK and XG analyzed the data, performed bioinformatic analysis, and drafted the manuscript. ZT and NS revised the final manuscript. All of the authors have approved the final version of the manuscript.

## Acknowledgments

Not applicable.

---

**Supplemental Figure 1.** mRNA expression of Il1-β, Tnf-α, and Il6 in mice lung tissues from the indicated group of mice in Figure 1 (A) was detected by qPCR (n = 8 per group). The results are shown as mean ± SD. Statistical testing was performed using an unpaired Student’s t-test. *p < 0.05, **p < 0.01, ***p < 0.001, ****p < 0.0001.

**Supplemental Figure 2.** Overview of infiltrating cell types in silica-induced lung fibrosis model. (A) t-SNE plot for single cells derived from lung tissues of mice with or without silicosis, color-coded by cell types. (B) t-SNE plot for single cells derived from lung tissues of mice with or without silicosis, color-coded by groups (n = 3 in silicosis mice group and n =2 in control mice group). (C) Dot plot depicting the average expression levels of canonical cell type-specific marker genes. The dot size shows the fraction of expressing cells, and the dots are colored based on the gene expression levels. (D) Bar plots showing the percent of different types of cells in total cells from each group.

**Supplemental Figure 3.** Dissection of myeloid cells indicating the potential pro-inflammatory and pro-fibrotic role of macrophages during silica exposure. (A) t-SNE plot for myeloid cells, color-coded by cell types. (B) t-SNE plot for myeloid cells, color-coded by groups (n = 3 in silicosis mice group and n =2 in control mice group). (C) Dot plot depicting each cell type’s average expression levels of canonical markers. The dot size shows the fraction of expressing cells, and the dots are colored based on the gene expression levels. (D) Bar plots showing the average proportion of each myeloid cell type from each group. (E) Heatmap of gene expression of pro-inflammatory genes (Il6, Il18, Il1b, Il1a, Tnf), selected differentially expressed chemokines detected in BALFs in Figure 1 (Ccl2, Ccl3, Ccl7, Cxcl1, Cxcl2, Cxcl5), and profibrotic genes (Il10, Mmp12, Igf1, Pdgfa, Pdgfb, Spp1, Tgfb1, Tgfbi, Lgmn, Mmp9, Mmp14, Timp1) for each myeloid cell subpopulation from all samples.

**Supplemental Figure 4.** (A) Differentiation trajectory inferred of Mmp12^hi^ macrophage via Monocle2 using Ly6c^+^ monocytes and all macrophages from all samples, colored by derived pseudotime (left) and cell type (right). (B) Heatmap of the area under the curve (AUC) scores of expression regulation by transcription factors, as estimated using pySCENIC. Shown are the top five differential transcription factors in each macrophage subtype.

**Supplemental Figure 5.** (A) t-SNE plot for fibroblasts derived from lung tissues of mice with or without silicosis, color-coded by cell types. (B) t-SNE plot for fibroblasts derived from lung tissues of mice with or without silicosis, color-coded by groups (n = 3 in silicosis mice group and n =2 in control mice group). (C) Dot plot depicting the average expression levels of canonical fibroblasts cell type-specific marker genes. (D) Bar plots showing the percent of different types of fibroblasts in total fibroblasts from each group (n = 3 in silicosis mice group and n =2 in control mice group). (E) Representative images of immunohistochemical staining of Cd68 (top), Col1a1 (middle), and Ly6g (bottom) expression in control (left) and silicosis (right) mice lung tissues (n = 5 per group, and scale bars=50 µm). (F) The GO pathway enrichment analysis using spatial DEGs of “nodules of lesion” versus “non-nodular area”.

Supplemental Figure 6. (A) t-SNE plot for single cells derived from BALFs of patients with or without silicosis, color-coded by cell types. (B) Dot plot depicting the average expression levels of canonical cell type-specific marker genes. (C) t-SNE plot for myeloid cells derived from BALFs of patients with or without silicosis, color-coded by cell types. (D) Dot plot depicting the average expression levels of canonical myeloid cells type-specific marker genes. (E) The GO pathway enrichment analysis using DEGs of Group 2 macrophages versus others to explore the functions of Group 2 macrophages.

