## Supplemental Figures for "Integrated multi-omics analyses reveal the pro-inflammatory and pro-fibrotic pulmonary macrophage subcluster in silicosis"

Supplemental Figure 1

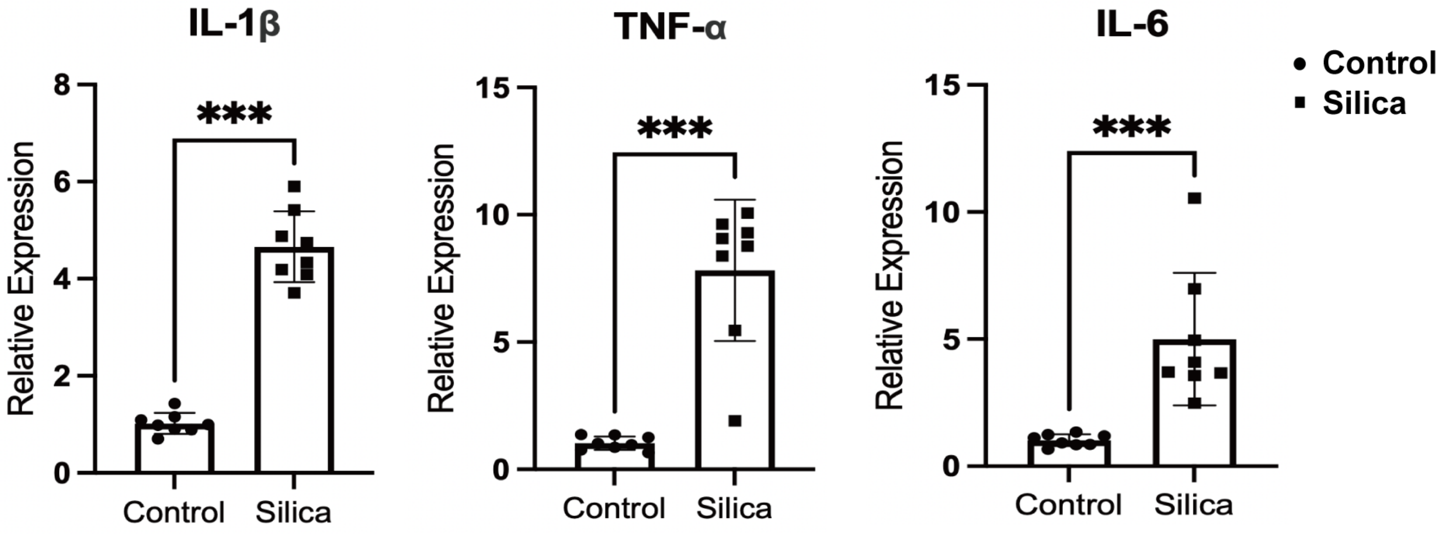

Supplemental Figure 2

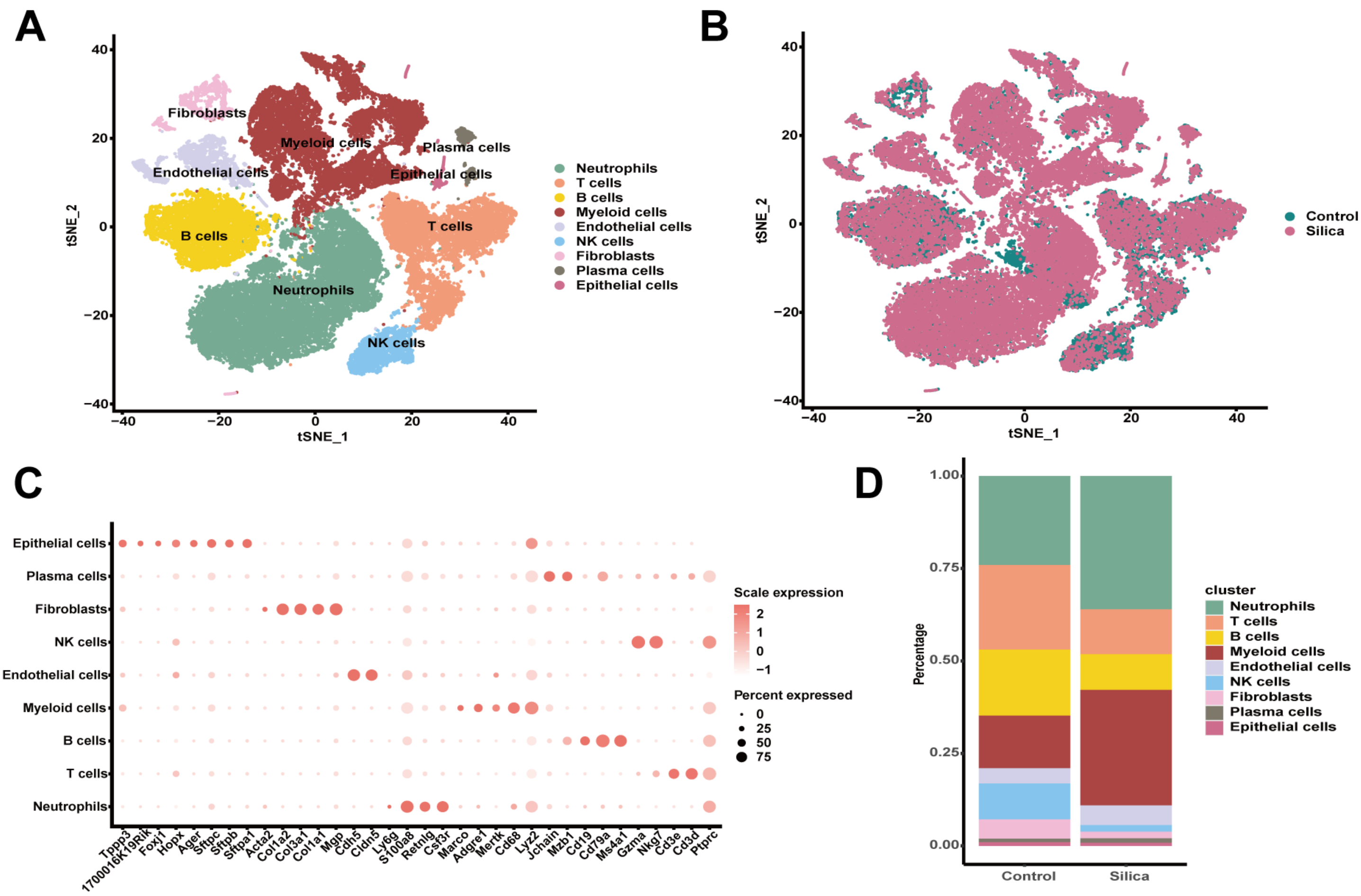

Supplemental Figure 3

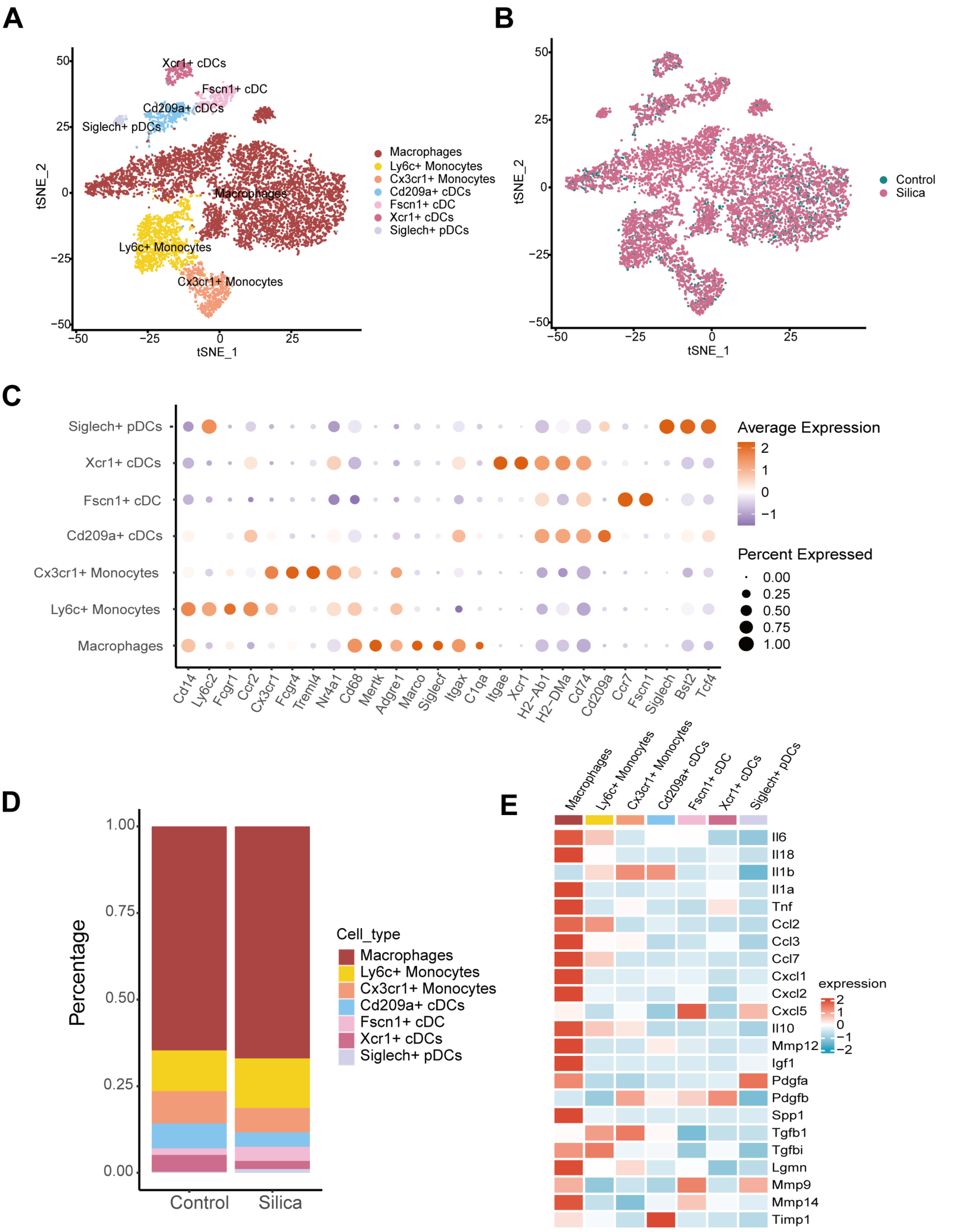

Supplemental Figure 4

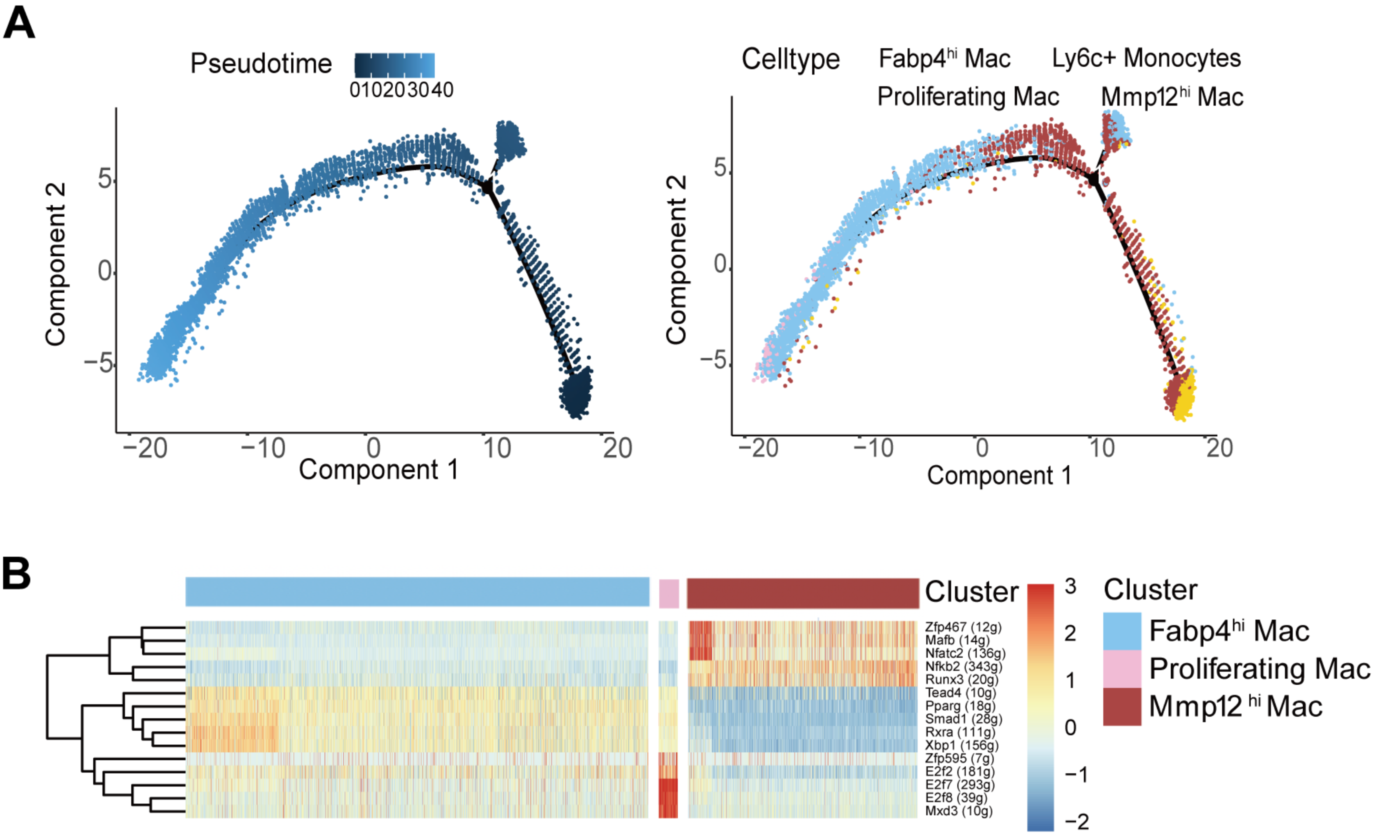

Supplemental Figure 5

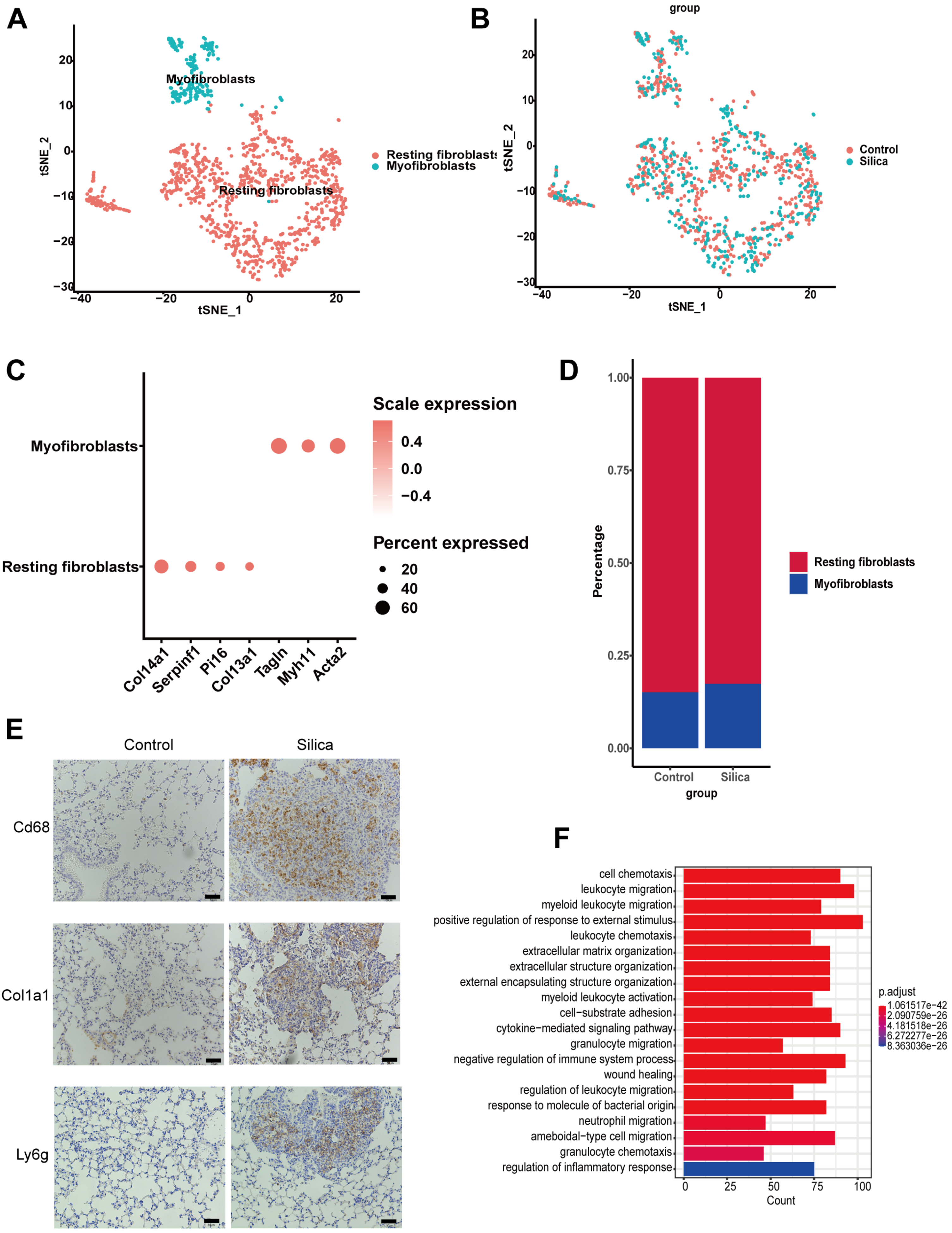

Supplemental Figure 6

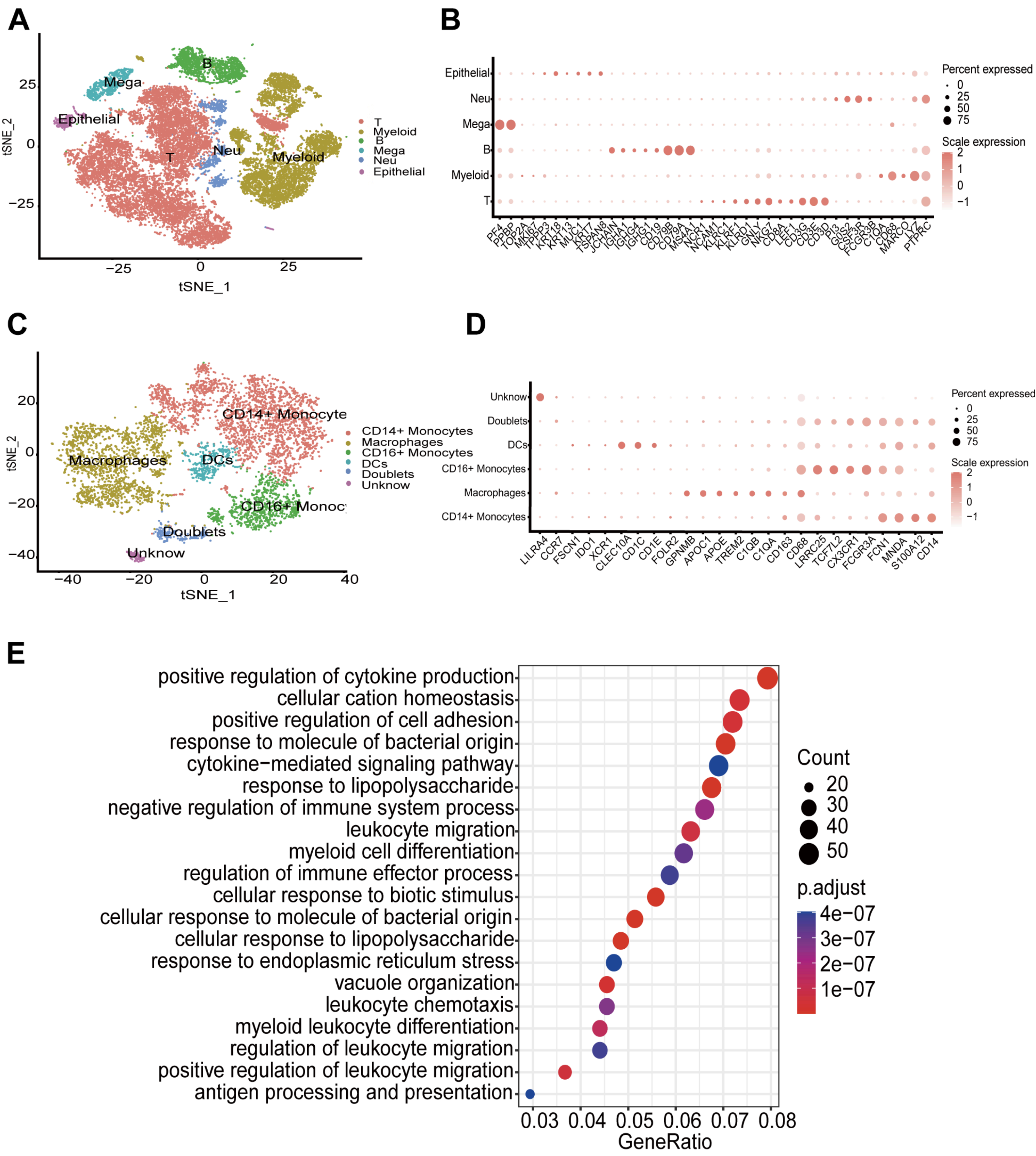
